## Supplemental Text for "Different structural variant prediction tools yield considerably different results in *Caenorhabditis elegans*"

Supplemental_Table_S1: Prediction of deletions from simulated short-read data

| | 5X | | | | 15X | | | | 30X | | | | 60X | | | |
|---|---|---|---|---|---|---|---|---|---|---|---|---|---|---|---|---|
| | **P** | **R** | **F1** | **J** | **P** | **R** | **F1** | **J** | **P** | **R** | **F1** | **J** | **P** | **R** | **F1** | **J** |
| BreakDancer | 0.93 | 0.76 | 0.84 | 0.83 | 0.96 | 0.89 | 0.92 | 0.95 | 0.95 | 0.88 | 0.92 | 0.95 | 0.94 | 0.88 | 0.91 | 0.95 |
| cnMOPS | 0.83 | 0.32 | 0.46 | 0.81 | 0.86 | 0.62 | 0.72 | 0.9 | 0.73 | 0.78 | 0.76 | 0.82 | 0.48 | 0.86 | 0.62 | 0.74 |
| CNVnator | **1.0** | 0.36 | 0.53 | 0.91 | **1.0** | 0.66 | 0.79 | 0.97 | 0.99 | 0.79 | 0.88 | 0.98 | 0.81 | 0.88 | 0.84 | **0.99** |
| Delly | **1.0** | **0.85** | **0.92** | **0.94** | **1.0** | **0.96** | **0.98** | **0.99** | **1.0** | **0.97** | **0.98** | **0.99** | **0.98** | **0.97** | **0.98** | 0.74 |
| Hydra | 0.93 | 0.66 | 0.78 | 0.12 | 0.89 | 0.71 | 0.79 | 0.13 | 0.85 | 0.71 | 0.77 | 0.12 | 0.79 | 0.67 | 0.72 | 0.12 |
| Lumpy | **1** | 0.76 | 0.87 | 0.78 | **1.0** | 0.93 | 0.96 | 0.96 | 0.99 | 0.93 | 0.96 | 0.96 | 0.41 | 0.94 | 0.57 | 0.95 |
| FusorSV[1] | 1.0 | 0.84 | 0.92 | 0.96 | 1.0 | 0.94 | 0.97 | 0.99 | 1.0 | 0.94 | 0.97 | 0.98 | 1.0 | 0.95 | 0.97 | 0.98 |

1. FusorSV used a training model trained on simulated data for the other callers.
2. P = precision, R = recall, F1 = F1 score, J = Jaccard similarity

Supplemental_Table_S2: Prediction of duplications from simulated short-read data

| | 5X | | | | 15X | | | | 30X | | | | 60X | | | |
|---|---|---|---|---|---|---|---|---|---|---|---|---|---|---|---|---|
| | P | R | F1 | J | P | R | F1 | J | P | R | F1 | J | P | R | F1 | J |
| BreakDancer | 0.99 | 0.75 | 0.85 | 0.77 | **1** | 0.86 | 0.93 | 0.88 | **1** | 0.86 | 0.93 | 0.88 | **1** | 0.86 | 0.93 | 0.88 |
| cnMOPS | 0.61 | 0.74 | 0.67 | 0.84 | 0.35 | 0.79 | 0.48 | 0.85 | 0.2 | 0.82 | 0.32 | 0.72 | 0.11 | 0.88 | 0.2 | 0.69 |
| CNVnator | **1** | 0.65 | 0.79 | **0.97** | 0.99 | 0.73 | 0.84 | **0.98** | 0.99 | 0.8 | 0.88 | **0.98** | 0.95 | 0.82 | 0.88 | **0.97** |
| Delly | **1** | **0.87** | **0.93** | 0.93 | **1** | **0.92** | **0.96** | 0.95 | 0.99 | **0.93** | **0.96** | 0.95 | 0.99 | **0.93** | **0.96** | 0.83 |
| Hydra | 0.36 | 0.26 | 0.30 | **0.01** | 0.56 | 0.28 | 0.37 | 0.01 | 0.77 | 0.27 | 0.4 | 0.01 | 0.77 | 0.26 | 0.38 | 0.01 |
| Lumpy | **1** | 0.75 | 0.86 | 0.77 | **1** | 0.87 | 0.93 | 0.87 | **1** | 0.9 | 0.95 | 0.9 | **1** | 0.91 | 0.95 | 0.92 |
| FusorSV[1] | 1.0 | 0.88 | 0.94 | 0.98 | 1.0 | 0.94 | 0.97 | 0.96 | 1.0 | 0.93 | 0.97 | 0.96 | 1.0 | 0.93 | 0.96 | 0.95 |

1. FusorSV used a training model trained on simulated data for the other callers.
2. P = precision, R = recall, F1 = F1 score, J = Jaccard similarity

Supplemental_Table_S3: Prediction of inversions from simulated short-read data

| | 5X | | | | 15X | | | | 30X | | | | 60X | | | |
|---|---|---|---|---|---|---|---|---|---|---|---|---|---|---|---|---|
| | P | R | F1 | J | P | R | F1 | J | P | R | F1 | J | P | R | F1 | J |
| BreakDancer | 0.95 | 0.9 | 0.92 | 0.94 | 0.93 | 0.91 | 0.92 | **0.95** | 0.93 | 0.91 | 0.92 | **0.95** | 0.93 | 0.92 | 0.92 | **0.96** |
| Delly | 0.96 | **0.94** | **0.95** | **0.97** | 0.94 | **0.95** | 0.95 | 0.78 | 0.94 | **0.96** | 0.95 | 0.78 | 0.93 | **0.95** | 0.94 | 0.77 |

| | | | | | | | | | | | | | | | | |
|---|---|---|---|---|---|---|---|---|---|---|---|---|---|---|---|---|
| Hydra | 0.98 | 0.28 | 0.43 | 0.32 | 0.4 | 0.02 | 0.04 | 0.0 | 0.27 | 0.02 | 0.04 | 0.01 | 0.21 | 0.01 | 0.03 | **0** |
| Lumpy | **1** | 0.9 | 0.94 | 0.89 | **1** | 0.93 | **0.97** | 0.92 | **1** | 0.94 | **0.97** | 0.94 | **1** | 0.94 | **0.97** | 0.94 |
| FusorSV[1] | 0.98 | 0.92 | 0.95 | 0.92 | 0.99 | 0.93 | 0.96 | 0.37 | 0.98 | 0.92 | 0.95 | 0.23 | 0.99 | 0.94 | 0.97 | 0.39 |

1. FusorSV used a training model trained on simulated data for the other callers.
2. P = precision, R = recall, F1 = F1 score, J = Jaccard similarity

Supplemental_Table_S4: Degree of overlap for deletions spanning genes predicted from long-read data and FusorSV

| Caller | Overlap with other callers | | | | | | | |
|---|---|---|---|---|---|---|---|---|
| | No others | One other | Two others | Three others | Four others | Five others | Total | Unique to caller (%) |
| assemblytics | 2998 | 206 | 66 | 81 | 104 | 119 | 3574 | 83.9 |
| mumandco | 409 | 210 | 121 | 170 | 231 | 119 | 1260 | 32.5 |
| pbsv | 374 | 184 | 198 | 229 | 230 | 119 | 1334 | 28.0 |
| sniffles | 1095 | 453 | 303 | 253 | 232 | 119 | 2455 | 44.6 |
| svim | 49 | 183 | 247 | 236 | 222 | 119 | 1056 | 4.60 |
| FusorSV | 1322 | 196 | 148 | 111 | 166 | 119 | 2062 | 64.1 |

Supplemental_Table_S5: Degree of overlap for deletions spanning genes predicted from short-read data

| Caller | Overlap with other callers | | | | | | | | |
|---|---|---|---|---|---|---|---|---|---|
| | No others | One other | Two others | Three others | Four others | Five others | Six others | Total | Unique to caller (%) |
| BreakDancer | 141 | 174 | 334 | 499 | 375 | 153 | 90 | 1766 | 8.0 |
| cnMOPs | 3847 | 450 | 86 | 63 | 115 | 131 | 90 | 4782 | 8.0 |
| CNVnator | 3368 | 558 | 49 | 23 | 46 | 72 | 90 | 4206 | 8.0 |
| Delly | 56 | 171 | 404 | 539 | 383 | 160 | 90 | 1803 | 3.1 |
| Hydra | 733 | 236 | 154 | 180 | 333 | 147 | 90 | 1873 | 39.1 |
| Lumpy | 81 | 179 | 389 | 533 | 389 | 156 | 90 | 1817 | 4.5 |
| FusorSV | 374 | 400 | 228 | 463 | 354 | 153 | 90 | 2062 | 18.1 |

Supplemental_Table_S6: Degree of overlap for duplications spanning genes predicted from long-read data and FusorSV

| Caller | Overlap with other callers | | | | | | | |
|---|---|---|---|---|---|---|---|---|
| | No others | One other | Two others | Three others | Four others | Five others | Total | Unique to caller (%) |
| assemblytics | 69 | 65 | 61 | 66 | 35 | 52 | 348 | 19.8 |
| mumandco | 25 | 31 | 29 | 54 | 29 | 52 | 220 | 11.4 |
| pbsv | 170 | 28 | 37 | 75 | 45 | 52 | 407 | 41.8 |
| sniffles | 816 | 565 | 244 | 114 | 46 | 52 | 1837 | 44.4 |
| svim | 145 | 448 | 247 | 117 | 47 | 52 | 1056 | 13.7 |
| FusorSV | 1428 | 145 | 156 | 46 | 33 | 52 | 1860 | 76.8 |

Supplemental_Table_S7: Degree of overlap for duplications spanning genes predicted from short-read data

| Caller | Overlap with other callers | | | | | | | | |
|---|---|---|---|---|---|---|---|---|---|
| | No others | One other | Two others | Three others | Four others | Five others | Six others | Total | Unique to caller (%) |
| BreakDancer | 77 | 30 | 58 | 236 | 110 | 15 | 1 | 527 | 14.6 |
| cnMOPs | 86 | 14 | 0 | 2 | 1 | 3 | 1 | 107 | 80.4 |
| CNVnator | 1386 | 81 | 13 | 21 | 29 | 16 | 1 | 1547 | 89.6 |
| Delly | 79 | 265 | 672 | 476 | 124 | 16 | 1 | 1633 | 4.8 |
| Hydra | 637 | 91 | 109 | 252 | 112 | 14 | 1 | 1216 | 52.4 |
| Lumpy | 109 | 220 | 624 | 474 | 124 | 16 | 1 | 1568 | 7.0 |
| FusorSV | 332 | 265 | 606 | 443 | 120 | 16 | 1 | 1783 | 18.6 |

Supplemental_Table_S8: Degree of overlap for inversions spanning genes predicted from long-read data and FusorSV

| Caller | Overlap with other callers | | | | | | |
|---|---|---|---|---|---|---|---|
| | No others | One other | Two others | Three others | Four others | Total | Unique to caller (%) |
| mumandco | 215 | 55 | 29 | 55 | 7 | 361 | 59.6 |
| pbsv | 222 | 103 | 44 | 61 | 7 | 437 | 50.8 |
| sniffles | 603 | 157 | 43 | 59 | 7 | 869 | 69.4 |
| svim | 115 | 77 | 24 | 60 | 7 | 283 | 40.6 |
| FusorSV | 601 | 46 | 13 | 9 | 7 | 676 | 88.9 |

Supplemental_Table_S9: Degree of overlap for inversions spanning genes predicted from short-read data

| Caller | Overlap with other callers | | | | | | |
|---|---|---|---|---|---|---|---|
| | No others | One other | Two others | Three others | Four others | Total | Unique to caller (%) |
| BreakDancer | 418 | 561 | 218 | 68 | 19 | 1284 | 32.6 |
| Delly | 411 | 606 | 238 | 70 | 19 | 1344 | 30.6 |
| Hydra | 200 | 74 | 81 | 56 | 19 | 430 | 46.5 |
| Lumpy | 21 | 30 | 30 | 26 | 19 | 126 | 16.7 |
| FusorSV | 218 | 191 | 180 | 68 | 19 | 676 | 32.2 |

Supplemental_Table_S10: Deletion performance for different SVIM QUAL cut-off thresholds

| Depth | Cut-off | Precision | Recall | F1 |
|---|---|---|---|---|
| 5X | 0 | 0.120 | 0.900 | 0.212 |
| 5X | 1 | 0.120 | 0.900 | 0.212 |
| 5X | 2 | 0.857 | 0.840 | 0.848 |
| 5X | 3 | 0.991 | 0.727 | 0.838 |
| 5X | 4 | 0.988 | 0.533 | 0.693 |
| 5X | 5 | 0.980 | 0.333 | 0.498 |
| 5X | 6 | 1.000 | 0.213 | 0.352 |
| 5X | 7 | 0.857 | 0.840 | 0.848 |
| 5X | 8 | 1.000 | 0.140 | 0.246 |
| 5X | 9 | 1.000 | 0.073 | 0.137 |
| 5X | 10 | 1.000 | 0.033 | 0.065 |
| 15X | 0 | 0.054 | 0.927 | 0.101 |
| 15X | 1 | 0.054 | 0.933 | 0.102 |
| 15X | 2 | 0.493 | 0.933 | 0.645 |
| 15X | 3 | 0.880 | 0.927 | 0.903 |
| 15X | 4 | 0.972 | 0.927 | 0.949 |
| 15X | 5 | 0.979 | 0.913 | 0.945 |
| 15X | 6 | 0.979 | 0.913 | 0.945 |
| 15X | 7 | 0.985 | 0.900 | 0.941 |
| 15X | 8 | 0.992 | 0.880 | 0.933 |
| 15X | 9 | 0.992 | 0.873 | 0.929 |
| 15X | 10 | 0.992 | 0.827 | 0.902 |
| 30X | 0 | 0.034 | 0.933 | 0.065 |
| 30X | 1 | 0.034 | 0.933 | 0.065 |
| 30X | 2 | 0.243 | 0.933 | 0.386 |
| 30X | 3 | 0.657 | 0.933 | 0.771 |
| 30X | 4 | 0.922 | 0.940 | 0.931 |
| 30X | 5 | 0.966 | 0.940 | 0.953 |
| 30X | 6 | 0.966 | 0.940 | 0.953 |

| | | | | |
|---|---|---|---|---|
| 30X | 7 | 0.979 | 0.940 | 0.959 |
| 30X | 8 | 0.986 | 0.940 | 0.962 |
| 30X | 9 | 0.993 | 0.933 | 0.962 |
| 30X | 10 | 0.993 | 0.927 | 0.959 |
| 60X | 10 | 0.979 | 0.933 | 0.956 |
| 60X | 11 | 0.986 | 0.940 | 0.962 |
| 60X | 12 | 0.979 | 0.933 | 0.956 |
| 60X | 13 | 0.986 | 0.940 | 0.962 |
| 60X | 14 | 0.986 | 0.940 | 0.962 |
| 60X | 15 | 0.979 | 0.933 | 0.956 |
| 142X | 15 | 0.972 | 0.940 | 0.956 |
| 142X | 20 | 0.986 | 0.940 | 0.962 |
| 142X | 25 | 0.986 | 0.940 | 0.962 |
| 142X | 30 | 0.993 | 0.933 | 0.962 |
| 142X | 35 | 0.993 | 0.933 | 0.962 |
| 142X | 40 | 0.993 | 0.927 | 0.959 |
| 142X | 45 | 0.993 | 0.927 | 0.959 |
| 142X | 50 | 0.993 | 0.927 | 0.959 |
| 142X | 55 | 0.993 | 0.920 | 0.955 |

Supplemental_Table_S11: Duplication performance for different SVIM QUAL cut-off thresholds

| Depth | Cut-off | Precision | Recall | F1 |
|---|---|---|---|---|
| 5X | 0 | 0.023 | 0.350 | 0.043 |
| 5X | 1 | 0.023 | 0.350 | 0.043 |
| 5X | 2 | 0.792 | 0.317 | 0.452 |
| 5X | 3 | 1.000 | 0.300 | 0.462 |
| 5X | 4 | 1.000 | 0.250 | 0.400 |
| 5X | 5 | 1.000 | 0.167 | 0.286 |
| 5X | 6 | 1.000 | 0.067 | 0.125 |
| 5X | 7 | 0.792 | 0.317 | 0.452 |
| 5X | 8 | 1.000 | 0.033 | 0.065 |
| 5X | 9 | 0 | 0 | 0 |
| 5X | 10 | 0 | 0 | 0 |
| 15X | 0 | 0.010 | 0.417 | 0.019 |
| 15X | 1 | 0.010 | 0.417 | 0.019 |
| 15X | 2 | 0.209 | 0.400 | 0.274 |
| 15X | 3 | 0.857 | 0.400 | 0.545 |
| 15X | 4 | 1.000 | 0.400 | 0.571 |
| 15X | 5 | 1.000 | 0.400 | 0.571 |
| 15X | 6 | 1.000 | 0.400 | 0.571 |
| 15X | 7 | 1.000 | 0.400 | 0.571 |
| 15X | 8 | 1.000 | 0.400 | 0.571 |
| 15X | 9 | 1.000 | 0.383 | 0.554 |
| 15X | 10 | 1.000 | 0.383 | 0.554 |

| | | | | |
|---|---|---|---|---|
| 30X | 0 | 0.006 | 0.450 | 0.012 |
| 30X | 1 | 0.006 | 0.450 | 0.012 |
| 30X | 2 | 0.082 | 0.433 | 0.138 |
| 30X | 3 | 0.400 | 0.433 | 0.416 |
| 30X | 4 | 0.897 | 0.433 | 0.584 |
| 30X | 5 | 0.963 | 0.433 | 0.598 |
| 30X | 6 | 0.963 | 0.433 | 0.598 |
| 30X | 7 | 1.000 | 0.433 | 0.605 |
| 30X | 8 | 1.000 | 0.433 | 0.605 |
| 30X | 9 | 1.000 | 0.433 | 0.605 |
| 30X | 10 | 1.000 | 0.433 | 0.605 |
| 60X | 10 | 1.000 | 0.467 | 0.636 |
| 60X | 11 | 1.000 | 0.467 | 0.636 |
| 60X | 12 | 1.000 | 0.467 | 0.636 |
| 60X | 13 | 1.000 | 0.467 | 0.636 |
| 60X | 14 | 1.000 | 0.467 | 0.636 |
| 60X | 15 | 1.000 | 0.467 | 0.636 |
| 142X | 15 | 0.972 | 0.940 | 0.956 |
| 142X | 20 | 0.986 | 0.940 | 0.962 |
| 142X | 25 | 0.986 | 0.940 | 0.962 |
| 142X | 30 | 0.993 | 0.933 | 0.962 |
| 142X | 35 | 0.993 | 0.933 | 0.962 |
| 142X | 40 | 0.993 | 0.927 | 0.959 |
| 142X | 45 | 0.993 | 0.927 | 0.959 |
| 142X | 50 | 0.993 | 0.927 | 0.959 |
| 142X | 55 | 0.993 | 0.92 | 0.955 |

Supplemental_Table_S12: Inversion performance for different SVIM QUAL cut-off thresholds

| Depth | Cut-off | Precision | Recall | F1 |
|---|---|---|---|---|
| 5X | 0 | 0.027 | 0.652 | 0.052 |
| 5X | 1 | 0.093 | 0.638 | 0.163 |
| 5X | 2 | 0.457 | 0.536 | 0.493 |
| 5X | 3 | 0.828 | 0.348 | 0.490 |
| 5X | 4 | 1.000 | 0.174 | 0.296 |
| 5X | 5 | 1.000 | 0.101 | 0.184 |
| 5X | 6 | 1.000 | 0.029 | 0.056 |
| 5X | 7 | 0.457 | 0.536 | 0.493 |
| 5X | 8 | 0 | 0 | 0 |
| 5X | 9 | 0 | 0 | 0 |
| 5X | 10 | 0 | 0 | 0 |
| 15X | 0 | 0.014 | 0.667 | 0.027 |
| 15X | 1 | 0.035 | 0.667 | 0.066 |
| 15X | 2 | 0.142 | 0.667 | 0.235 |
| 15X | 3 | 0.338 | 0.667 | 0.449 |

| | | | | |
|---|---|---|---|---|
| 15X | 4 | 0.590 | 0.667 | 0.626 |
| 15X | 5 | 0.836 | 0.667 | 0.742 |
| 15X | 6 | 0.836 | 0.667 | 0.742 |
| 15X | 7 | 0.978 | 0.652 | 0.783 |
| 15X | 8 | 0.976 | 0.594 | 0.739 |
| 15X | 9 | 1.000 | 0.565 | 0.722 |
| 15X | 10 | 1.000 | 0.493 | 0.660 |
| 30X | 0 | 0.010 | 0.667 | 0.020 |
| 30X | 1 | 0.022 | 0.667 | 0.042 |
| 30X | 2 | 0.058 | 0.667 | 0.107 |
| 30X | 3 | 0.126 | 0.667 | 0.212 |
| 30X | 4 | 0.231 | 0.667 | 0.343 |
| 30X | 5 | 0.426 | 0.667 | 0.520 |
| 30X | 6 | 0.426 | 0.667 | 0.520 |
| 30X | 7 | 0.592 | 0.652 | 0.621 |
| 30X | 8 | 0.703 | 0.652 | 0.677 |
| 30X | 9 | 0.833 | 0.652 | 0.732 |
| 30X | 10 | 0.918 | 0.652 | 0.763 |
| 60X | 10 | 0.441 | 0.652 | 0.526 |
| 60X | 11 | 0.441 | 0.652 | 0.526 |
| 60X | 12 | 0.549 | 0.652 | 0.596 |
| 60X | 13 | 0.634 | 0.652 | 0.643 |
| 60X | 14 | 0.789 | 0.652 | 0.714 |
| 60X | 15 | 0.833 | 0.652 | 0.732 |
| 142X | 15 | 0.153 | 0.652 | 0.248 |
| 142X | 20 | 0.304 | 0.652 | 0.415 |
| 142X | 25 | 0.511 | 0.652 | 0.573 |
| 142X | 30 | 0.918 | 0.652 | 0.763 |
| 142X | 35 | 1.000 | 0.652 | 0.789 |
| 142X | 40 | 1.000 | 0.638 | 0.779 |
| 142X | 45 | 1.000 | 0.638 | 0.779 |
| 142X | 50 | 1.000 | 0.623 | 0.768 |
| 142X | 55 | 1.000 | 0.449 | 0.62 |

Supplemental_Table_S13: Criteria used to select best overlapping call

| Caller | Criteria used to select best call | Notes |
|---|---|---|
| BreakDancer | QUAL | QUAL is the Phred-scale quality score for the assertion made in ALT 10log (prob(call in ALT is wrong)) |
| cnMOPs | N/A | Did not generate overlapping calls |
| CNVnator | N/A | Did not generate overlapping calls |
| DELLY | Max (paired_end_support, | Maximum read support from |

| | | |
|---|---|---|
| | split_read_support) | either paired-end or split reads |
| Hydra | None | |
| Lumpy | SU | Description="Number of pieces of evidence supporting the variant across all samples"> |
| Assemblytics | N/A | No pertinent information provided |
| MUM&Co | N/A | No pertinent information provided |
| pbsv | Read depth per allele | Read support for the variant allele |
| Sniffles | "# high-quality variant reads" | Read support for the variant allele |
| SVIM | svim qual score | Quality score specific to SVIM |
